# Therapeutic acute intermittent hypoxia does not alter whole-brain cerebrovascular function in healthy individuals

**DOI:** 10.64898/2026.10.06.757062

**Authors:** Hannah R Johnson, Max C Wang, Rachael C Stickland, Yufen Jennie Chen, Todd Parrish, Farzaneh A. Sorond, Molly G Bright

## Abstract

Acute intermittent hypoxia (AIH) therapy is a promising intervention known to elicit beneficial neuroplasticity. It has been suggested that AIH may also modulate cerebrovascular function, yet the cerebrovascular effects of specific therapeutic intermittent hypoxia protocols remain poorly characterized. Understanding whether AIH results in favorable or adverse cerebrovascular changes is critical in establishing the safety of AIH treatments in populations with elevated cerebrovascular risk. This study investigated cerebral blood flow, cerebrovascular reactivity to hypoxia, and blood pressure effects in a sham-controlled crossover trial (NCT05164705). Healthy young adults (n=18) completed three-week arms of AIH (12 sessions, 15 cycles of 60 seconds hypoxia alternated with 60 seconds normoxia) and sham (6 sessions, normoxia) treatments; phase-contrast and arterial spin labeling magnetic resonance imaging were used to quantify whole-brain and regional cerebral blood flow and hypoxic cerebrovascular reactivity. Blood pressure was measured acutely (before and after each intervention session) and longitudinally (before and after 3 weeks of treatment). AIH treatment did not significantly alter whole-brain resting cerebral blood flow or cerebrovascular reactivity to hypoxia. Blood pressure was unaffected by the treatment acutely or longitudinally. Voxel-wise analysis identified modest regional perfusion increases in the frontal pole, postcentral gyri, and precentral gyri with AIH relative to sham. Significant variability between subjects and between sessions in the hypoxic burden delivered with this AIH protocol was also identified. Overall, this three-week therapeutic AIH protocol produced minimal sustained hemodynamic changes in healthy young adults. The insights from this trial lay a foundation for safely expanding access to this promising rehabilitative intervention in clinical populations.

## 1 Introduction

Acute intermittent hypoxia (AIH) therapy has emerged as a promising rehabilitative treatment that improves motor function. Research in individuals with chronic, incomplete spinal cord injury has found that somatic motor function is enhanced by AIH treatment, with increased walking speed and endurance, hand function and dexterity, ventilation, and maximal inspiratory pressure reported^1–4^. The brief repeated exposures to hypoxia in AIH treatment increase motor output by activating serotonergic raphe nuclei, triggering a serotonin- and BDNF-dependent signaling cascade that ultimately augments synaptic strength and long-term excitability in cranial and spinal motoneuron pools^5–7^. This enhanced excitability leverages spared neural pathways in incomplete spinal cord injury, magnifying their synaptic signaling to motoneurons below the site of injury and ultimately increasing motor output^6^. Preliminary work has also suggested that AIH therapy may be a useful treatment in further clinical conditions where neural signaling is impaired, including Alzheimer’s disease, stroke, and multiple sclerosis^8,9^.

However, the physiological effects of hypoxia extend beyond inciting neuroplasticity. Hypoxia itself may directly injure tissues, with the brain being especially susceptible to hypoxic injury due to its high metabolic demand. Exposure to intermittent hypoxia can produce additional and extensive negative effects depending on the frequency and severity of hypoxic exposure, particularly in the cardiovascular system, as evidenced by obstructive sleep apnea^10^. Alternatively, therapeutic AIH may also positively impact cardio- and cerebrovascular function. Research has shown that hypoxic preconditioning, or exposure to subnoxious hypoxia, can improve tolerance to future exposures and protect against hypoxic injury by promoting cell survival and angiogenesis through hypoxia-inducible factor-1α upregulation, improving outcomes in stroke, COVID-19, and other cardiovascular and respiratory disorders^11,12^. Before this emerging therapeutic treatment can be safely applied in clinical populations that may be at greater risk for hypoxic damage, the cerebrovascular impacts of AIH therapy need to be assessed.

Cerebral blood flow (CBF) is significantly reduced with age and neurodegeneration as cerebrovascular network density decreases and its functional integrity diminishes^13–15^. Such decreases in CBF are associated with, and may in fact cause, cognitive impairment and increased risk of white matter lesions^16–19^. Aging and neurodegeneration are also associated with reduced cerebrovascular reactivity (CVR), or the diminished capacity of the cerebrovasculature to respond to alterations in cerebral perfusion pressure, neurocognitive demand, and nutrient availability, increasing risk of cerebrovascular accidents^20–22^. However, patient populations may also have existing vascular dysregulation; further impairments in autoregulation could put subjects at risk of cerebrovascular accidents and disease, as seen in obstructive sleep apnea^23^. Identifying the effect of therapeutic AIH protocols on cerebrovascular function is critical to ascertaining in what populations this rehabilitative treatment can be safely and effectively applied.

A handful of previous studies have assessed cerebrovascular function following intermittent hypoxia as an outcome measure, but the results of these studies have been mixed. Some have reported that CBF and CVR to hypoxia (HX-CVR) increased^24–28^, whereas other studies have reported that intermittent hypoxia did not change or even decreased CBF and HX-CVR^23,29^. Studied primarily in healthy adults, the disparity in reported effects of intermittent hypoxia exposure on cerebrovascular function is likely due, in part, to differences in the AIH protocol implemented by each study. It has been established that whether favorable or pathogenic outcomes result from intermittent hypoxia depends largely on the hypoxic dose delivered^9^. The intermittent hypoxia dose is dependent on the degree of hypoxia, the duration of each hypoxic exposure, the number of hypoxic exposures per session, the number of sessions per day, and the temporal distribution of sessions. Severe intermittent hypoxia doses are associated with a larger sympathetic response and greater production of reactive oxygen species than mild AIH doses, and they additionally activate microglia to a pro-inflammatory state, ultimately constraining the mechanisms that produce beneficial synaptic plasticity through cross-talk inhibition^30,31^. Dose comparison studies have shown that the degree of neuroprotection, strength enhancements, and cardiovascular improvements all vary with the hypoxic dose^6,9,32–35^. The same is likely true for cerebrovascular improvements.

Hypoxia is a hormetic stimulus of cerebrovascular function, with CBF initially increasing with mild, short-term exposures but with prolonged exposures resulting in injury and reduced reactivity to hypoxia and other cerebrovascular stimuli^36^. If intermittent hypoxia protocols reduce CBF or CVR, this could exacerbate the risk of cerebrovascular accidents and further impair cognitive function. If intermittent protocols instead *enhance* CBF and CVR, this could indicate strong therapeutic potential for improving cognition and neuroplasticity and for attenuating disease processes in aging, chronic stroke, chronic traumatic brain injury, vascular dementia, and psychiatric conditions, especially when other interventions such as aerobic exercise are not feasible^37–43^. At the same time, chronic elevations in CBF and CVR may be dangerous for individuals with acute stroke and acute traumatic brain injury or conditions with fragile cerebrovasculature such as Moyamoya disease, where impaired autoregulation and weak vessels may be unable to withstand changes in cerebral blood pressure^44,45^. As the cerebrovascular effects of therapeutic AIH are dose-dependent, it is critical to assess specific protocols for their outcomes.

We therefore sought to characterize the effects of an established rehabilitative AIH protocol on blood pressure, CBF, and HX-CVR. This protocol is already employed in treating spinal cord injury patients and is under investigation for further clinical applications in stroke and multiple sclerosis^46,47^.

Many methods exist to assess cerebrovascular function, including transcranial doppler ultrasound, positron emission tomography, near-infrared spectroscopy, and magnetic resonance imaging (MRI) approaches of phase contrast (PC), arterial spin labeling (ASL), dynamic susceptibility contrast, and blood oxygen level dependent (BOLD) imaging. PC, ASL, and BOLD MRI present significant advantages over other modalities as they are non-invasive, require no exogenous contrast agents, and can characterize whole-brain cerebrovascular dynamics. However, the BOLD signal is not quantitative and is confounded by bulk changes in blood oxygenation, complicating measures of CBF and HX-CVR, respectively^48^. PC and ASL MRI are more appropriate tools for both measures. PC MRI of the cerebral feeding arteries allows quantitative assessment of CBF free of the confounds encountered with BOLD MRI, but it is limited to whole-brain values. ASL MRI permits measurement of regional CBF and CVR, although it can be impacted by changes in labeling efficiency and arterial oxygenation during hypoxia^49^. Together, the complementary metrics of PC and ASL MRI can provide a comprehensive assessment of the impact of AIH therapy on cerebrovascular function.

Thus, a randomized, single-blind, sham-controlled crossover study was conducted to investigate the impact of a therapeutic AIH protocol on healthy cerebrovascular function measured with magnetic resonance imaging (MRI), using PC and ASL MRI to quantify whole-brain and regional CBF at rest. We also evaluated changes in CBF during a sustained hypoxic stimulus to quantify HX-CVR.

## 2 Methods

### 2.1 Study Design

This study (NCT05164705) was approved by the Institutional Review Board at Northwestern University, and all participants provided written informed consent before undergoing any study procedures. Eligible participants were between 21 and 50 years old, free of any known neurological, respiratory, or vascular conditions, safe to be scanned with MRI, not considered to be a frequent smoker (≤5 cigarettes/day), and had normal blood pressure at screening (90/50 mmHg < BP < 140/90 mmHg).

Participants attended a screening visit and were given the opportunity to practice gas inhalation under the AIH intervention and sustained gas delivery for MRI scan sessions to ensure comfort. Subjects underwent three weeks of AIH treatment or sham treatment, with MRI acquisitions performed before and after the intervention period. This protocol was repeated with the second treatment after a washout period of 3–5 months. Treatment order was randomized. Overall study design is depicted in Figure 1A.

**Figure 1.**
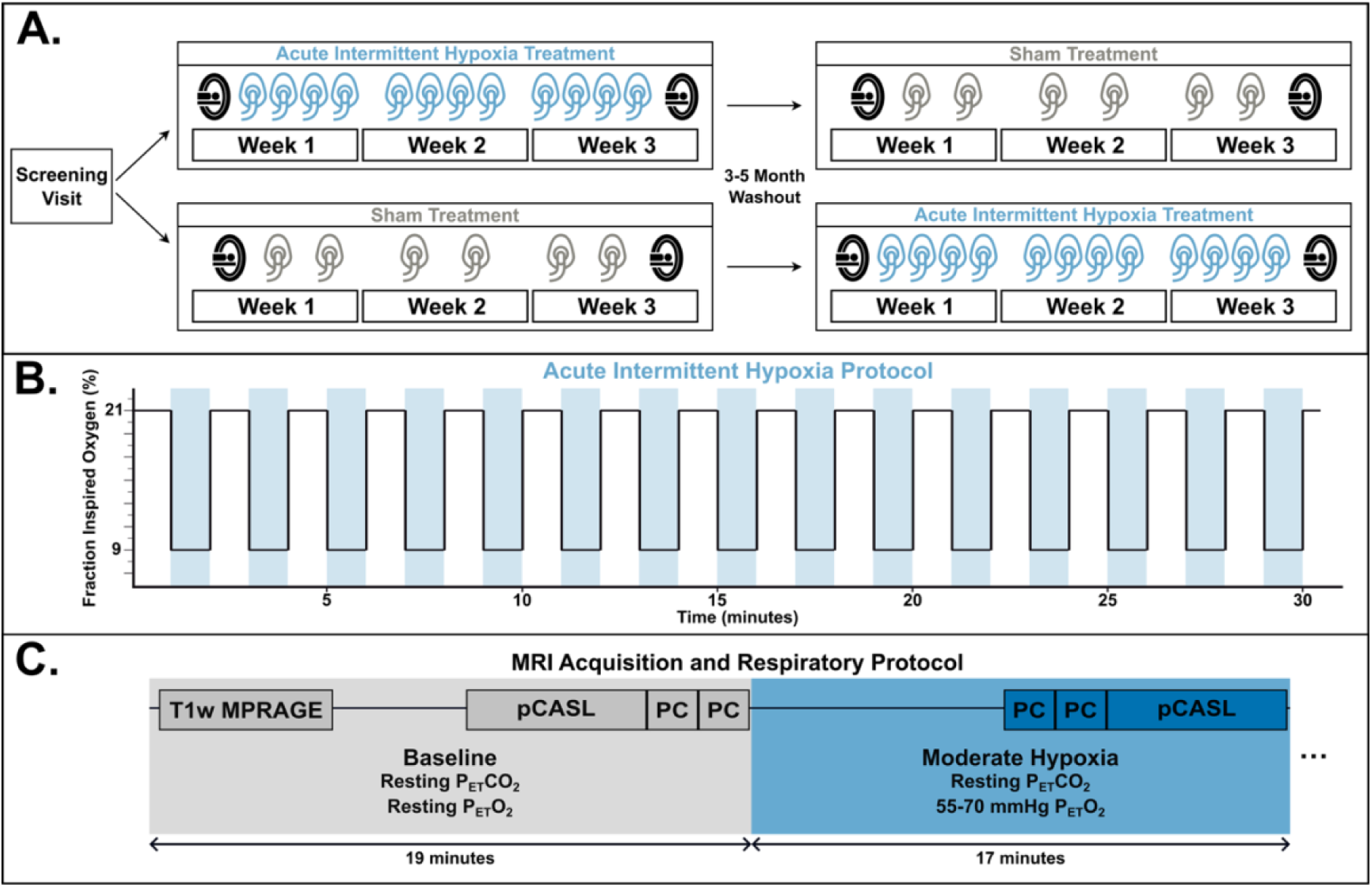
Study Design. Overall study design is shown in Panel A; participants underwent two three-week treatment arms, separated by a washout period of 3-5 months. In each arm, an MRI was performed before any intervention and another after all intervention sessions were completed. Acute intermittent hypoxia or sham treatment was given on four or two days, respectively, per week for the three weeks between MRI sessions. The target respiratory protocol for acute intermittent hypoxia treatment is shown in Panel B; blue shading indicates where participants inhaled hypoxic (9% O2) air from an altitude generator. The sham treatment respiratory protocol was identical except that the altitude generator output normoxic (21% O2) air. The MRI acquisition and respiratory protocol is shown in Panel C. The baseline respiratory period, lasting up to 19 minutes, maintained end-tidal gases at resting levels; the moderate hypoxia period, lasting up to 17 minutes, maintained end-tidal carbon dioxide at participants’ resting levels but targeted end-tidal oxygen levels between 55 and 70mmHg, depending on participants’ tolerance as determined in the screening visit. Actual timing of respiratory periods depended on the time required to collect each phase contrast acquisition.

### 2.2 AIH and Sham Intervention Protocols

In the AIH treatment arm, intervention sessions were held four times per week for a total of 12 sessions; in the sham treatment arm, intervention sessions were held twice per week for a total of six sessions to reduce participant burden. Participants were blinded to the oxygen content of the delivered gases but were told that the dose was being varied to account for the differing number of sessions. For all intervention sessions, participants were seated and fit with a non-rebreathing mask. Pulse, peripheral oxygen saturation (SpO_2_), end-tidal oxygen and carbon dioxide, and respiration were monitored and recorded at 100 Hz (Nonin Model 7500, Nonin Medical Inc, Plymouth, MN, USA; PowerLab 8/35 and Gas Analyzer, ADInstruments, Dunedin, New Zealand). Blood pressure measurements (Model BP7100, Omron Healthcare, Kyoto, Japan) were taken from the right arm immediately before and after the AIH or sham protocol while subjects were seated with arms supported at the level of the heart; three consecutive measurements were taken. Upon arrival to intervention sessions, participants sat and rested for five minutes before blood pressure measurements were taken, and one minute of rest was given between consecutive measurements.

The gas delivery protocol for both interventions lasted 30 minutes, consisting of 15 2-minute cycles (Fig. 1B). During each trial, hypoxia (9% FiO_2_) or sham hypoxia (21% FiO_2_) were delivered by connecting an altitude generator (Hyp-123, Hypoxico Inc, Gardiner, NY, USA) to the non-rebreathing face mask and were delivered for a period of one minute or until the participant’s SpO_2_ reached 85%, whichever threshold was met first. For the remainder of the 2-minute cycle, participants breathed room air (21% FiO_2_) to allow SpO_2_ to recover to resting levels.

### 2.3 MRI Protocol

Participants were scanned before and after the three-week intervention periods to measure CBF and CVR to moderate hypoxia (HX-CVR). A computer-controlled gas blending system (RespirAct, Thornhill Medical, Toronto, Canada) connected to an air-tight face mask was used to manipulate inhaled gas concentrations. Individual end-tidal partial pressures of oxygen (P_ET_O_2_) and carbon dioxide (P_ET_CO_2_) were prospectively targeted to create baseline, moderate hypoxic, and mild hypercapnic conditions during scanning. The gas protocol during scanning consisted of 4 phases: 1) up to 19 minutes of baseline, targeting individuals’ resting P_ET_O_2_ and P_ET_CO_2_ levels, 2) up to 17 minutes of moderate hypoxia, targeting resting P_ET_CO_2_ levels and P_ET_O_2_ of 55-70mmHg, 3) 5 minutes of rest, targeting individuals’ resting P_ET_O_2_ and P_ET_CO_2_ and allowing the effects of the moderate hypoxia condition to wash out, and 4) up to 14 minutes of mild hypercapnia, targeting resting P_ET_O_2_ levels and a P_ET_CO_2_ of 6mmHg higher than the individual’s baseline P_ET_CO_2_. Phases 3 and 4 were included to assess additional exploratory study endpoints. Only the initial baseline and hypoxia phases are examined in the current study. The respiratory phases of interest and corresponding MRI acquisitions are shown in Figure 1C.

Scanning was performed on a Siemens 3T Prisma scanner with a 32-channel head coil. Under baseline and moderate hypoxic respiratory conditions, two consecutive, retrospectively gated PC acquisitions (TR = 47.52ms, TE = 3.45ms, voxel size = 0.5×0.5×3mm^3^, encoding velocity = 100cm/s) were collected. Quantitative multi-post label delay (multi-PLD) pseudo-continuous arterial spin labeling (pCASL) acquisitions with background suppression (TR = 4100ms, TE = 30.56ms, voxel size = 3.5×3.5×3.5mm^3^, label duration=1.51s, post-label delays = 0.4/0.8/1.2/1.6/2.0s, repetitions = 2/2/2/3/3) were also obtained under each respiratory phase^50^. A T1-weighted MPRAGE (TR = 2170.0ms, TE = 1.69/3.55/5.41ms, voxel size = 1.0×1.0×1.0mm^3^) was acquired for registration and volume calculations, and a 3D time-of-flight angiogram (TR = 21.0ms, TE = 3.41ms, voxel size = 0.8×0.8×1.2mm^3^) was acquired at the level of the carotid bifurcation to position PC acquisitions and the labeling plane for pCASL acquisitions; both were positioned to maximize orthogonality to the internal carotid and vertebral arteries, just superior to the carotid bifurcation. Screenshots of the slice placement used in the first scan session were used to replicate placement in subsequent sessions.

### 2.4 Data and Statistical Analysis

Blood pressure measurements for each subject, session, and timepoint were averaged across the three repeated measures. SpO_2_ recordings were lowpass filtered (MATLAB lowpass, cutoff frequency = 0.5Hz) to produce a “desaturation curve”. The overall hypoxic burden for each AIH session was defined as the cumulative area between the desaturation curve and a 90% threshold^51,52^.

Image pre-processing for structural and ASL acquisitions was performed with the FMRIB Software Library (version 6.0). T1-weighted images were brain-extracted and segmented by tissue type (FSL_ANAT). Gray matter and white matter volumes were summed to calculate total brain volume for each participant, and brain mass was determined assuming tissue density of 1.06 g/mL^53,54^.

PC acquisitions were processed with cvi42 Flow 2D (Circle Cardiovascular Imaging): background phase correction was applied, regions of interest for the internal carotid and vertebral arteries were selected across each cardiac bin, and mean flow, calculated as the product of the average velocity and the cross-sectional area, was found for each artery. Total CBF from each phase contrast acquisition was calculated as the sum of mean flows for each of the four arteries. To align with perfusion measurements, normalized CBF in units of mL/100 g/min was found by dividing total CBF by the subject’s brain mass. Final normalized CBF measurements for the baseline and hypoxia phase of the MRI experiment were calculated as the average of the two consecutive acquisitions.

pCASL images were registered to the M0 image to correct for motion. Label-control image pairs were subtracted to isolate the perfusion signal and difference images for each post-label delay were averaged across repetitions. Quantitative maps of perfusion and other parameters were generated with FSL’s BASIL toolbox, using partial volume correction and assuming a labeling efficiency of 0.8^55–57^. Regional perfusion was analyzed by parcellating maps by vascular territory and finding the average perfusion for the right and left anterior, middle, and posterior cerebral artery regions^58,59^.

End-tidal data were temporally aligned with image acquisitions using start markers, which were placed synchronous with the start of the first baseline phase contrast acquisition during data collection, and average P_ET_O_2_ during each scan acquisition was calculated. HX-CVR to the moderate hypoxic stimulus for both pCASL and phase contrast data was calculated as the relative change in CBF modulated by the change in P_ET_O_2_ (Equation 1).

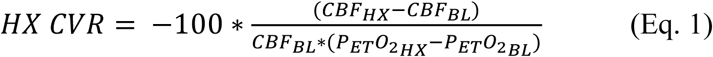

We quantified the hypoxic burden for each AIH intervention, calculated as the area spent under 90% SpO_2_ on a desaturation curve, plotting SpO_2_ over time^52^. Linear mixed effects models were used to assess differences in hypoxic burden; hypoxic burden was modeled with fixed effects of subject, session, bout, and the interaction between session and bout and a random intercept for each subject-session combination to account for nesting of bouts within sessions. We also employed linear mixed effects models to assess the intervention effect on cerebrovascular function, with subject included as a random effect and time and experimental arm as fixed effects. Coefficients of variation were calculated based on these models, with PC models for assessing variability additionally including the random effect of measurement repetition number. ASL measures of CBF were assessed on whole-brain, regional, and voxel-wise bases: regional analyses were conducted with the same linear mixed effects models, additionally including region as a fixed effect, and voxel-wise analyses were conducted with a three-factor ANOVA, including time and experimental arm as fixed effects and subject as a random effect. Additionally, the effect of cumulative hypoxic burden on cerebrovascular function was assessed with linear mixed effect models including subject as a fixed effect. Bland-Altman analysis, Pearson’s correlation coefficient, and intraclass correlation coefficient were used to assess agreement between PC and ASL CBF measures. Post-hoc pairwise comparisons with estimated marginal means were conducted and reported for blood pressure, CBF, and CVR linear mixed effects models. The interaction of time and experimental arm was used to test for overall significant differences associated with the AIH intervention. The Wilcoxon signed-rank test was used to compare blood pressure pre- and post-intervention sessions and between the first and last intervention session of each arm. A significance threshold of α=0.05 was used.

## 3 Results

A total of thirty-three individuals enrolled in the study. Eight participants were excluded following the initial screening visit after not meeting inclusion criteria (n=2) or declining further participation (n=6). A further seven participants did not complete all study activities due to scheduling conflicts (n=3), incidental imaging findings (n=1), reaching study stopping criteria during scanning (n=1), an anticipated side effect of anxiety during the intervention (n=1), and an unanticipated side effect of akathisia (n=1), which was deemed by physicians likely related to a new medication the participant initiated during the trial. A total of eighteen participants (9 female, 8 male, 1 gender-neutral, 29.0 ± 7.2 years) completed the study.

### 3.1 Effects of Gas Inhalation Protocols

During AIH interventions, the average minimum SpO_2_ reached in each hypoxic exposure was 85.13 ± 6.09%. An example desaturation curve recorded during an AIH intervention is shown in Figure 2A, with hypoxic burden highlighted; cumulative hypoxic burden for each subject and session is shown in Figure 2B. Subject, session, and bout number all significantly influenced the hypoxic burden delivered (p<0.001). The hypoxic burden increased by 2.36 ± 0.63 %×s with session and by 3.07 ± 0.30 %×s with bout. The hypoxic burden received was highly variable between subjects; subject-level fixed effects ranging from −97.42 to 66.49 %×s and between-subject variance accounted for over a third of total outcome variance.

**Figure 2.**
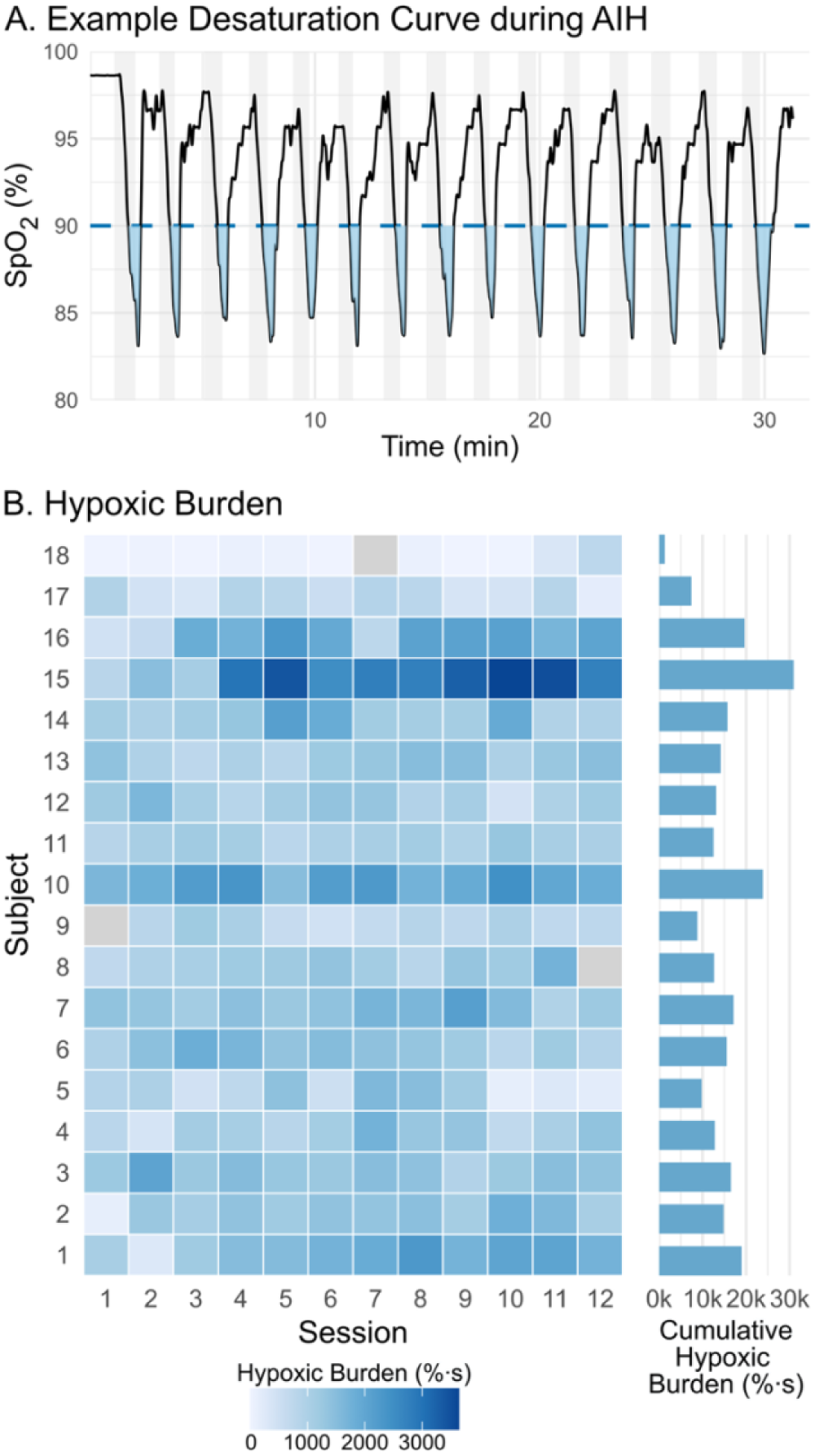
Acute Intermittent Hypoxia Intervention. A) An example desaturation curve recorded across the course of an AIH intervention, with the area under a 90% SpO2 threshold, indicated with a dashed line and shaded in blue; this area was used to calculate the hypoxic burden. Periods of hypoxic inhalation are shown in grey. B) Calculated hypoxic burden for each subject and session are shown in the heatmap on the left; missed (subject 8 session 12 and subject 18 session 7) and unrecorded (subject 9 session 1) sessions are indicated in grey. Cumulative hypoxic burden across all AIH sessions is shown for each subject on the right.

During baseline scanning, P_ET_CO_2_ and P_ET_O_2_ were maintained at 40.6±2.6 mmHg and 111.3±7.3 mmHg, respectively, an average of 0.9 and 1.2 mmHg, respectively, above the targeted levels. During the moderate hypoxia phase, P_ET_O_2_ averaged 62.8±4.8 mmHg, deviating an average of 1.6 mmHg from targeted levels, and P_ET_CO_2_ averaged 40.4±2.6 mmHg, deviating 0.6 mmHg on average from targeted values.

### 3.2 Quality, Repeatability, and Agreement of CBF Measurements

PC and pCASL images were visually inspected to verify data quality. PC data quality for all subjects who completed the study was satisfactory. Data quality in 15 pCASL datasets from 5 participants was qualitatively deemed insufficient based on the contrast between gray and white matter and gross asymmetry in perfusion signal, potentially related to poor labeling of inflowing arterial blood; these datasets were excluded from analyses. Coefficients of variation for PC measurements of CBF (considering only pre-intervention measures, n = 72), capturing total variation, variation between subjects, variation within subjects between scan sessions, and variation between scan repeats within a scan session, were 14%, 7.6%, 11%, and 4.4%, respectively, at baseline and 15.5%, 8.6%, 12.2%, and 4.4%, respectively, during moderate hypoxia. For ASL measurements in pre-intervention sessions (n = 28), coefficients of variation between subjects, within subjects between scan sessions, and in total were 11.7%, 15.4%, and 19.3%, respectively, at baseline and 15.1%, 12.1%, and 19.4% during hypoxia.

CBF values showed poor agreement but moderate correlations between PC and ASL measurements during both baseline and moderate hypoxia, seen in Figure 3. At baseline, the intraclass correlation coefficient between PC and ASL measurements of CBF was 0.55 and the Pearson’s correlation coefficient was 0.61. Mean bias at baseline was 4.0 mL/100g/min (95% limits of agreement: −13.5 to 21.5 mL/100g/min), with a proportional bias slope of −0.3 (p=0.03). During hypoxia, the intraclass correlation coefficient between the two modalities was 0.44 and the Pearson’s correlation coefficient was 0.55. Mean bias during hypoxia was 7.1 mL/100g/min (95% limits of agreement: −12.0 to 26.2 mL/100g/min) and the slope representing proportional bias was −0.15 (p=0.32).

**Figure 3.**
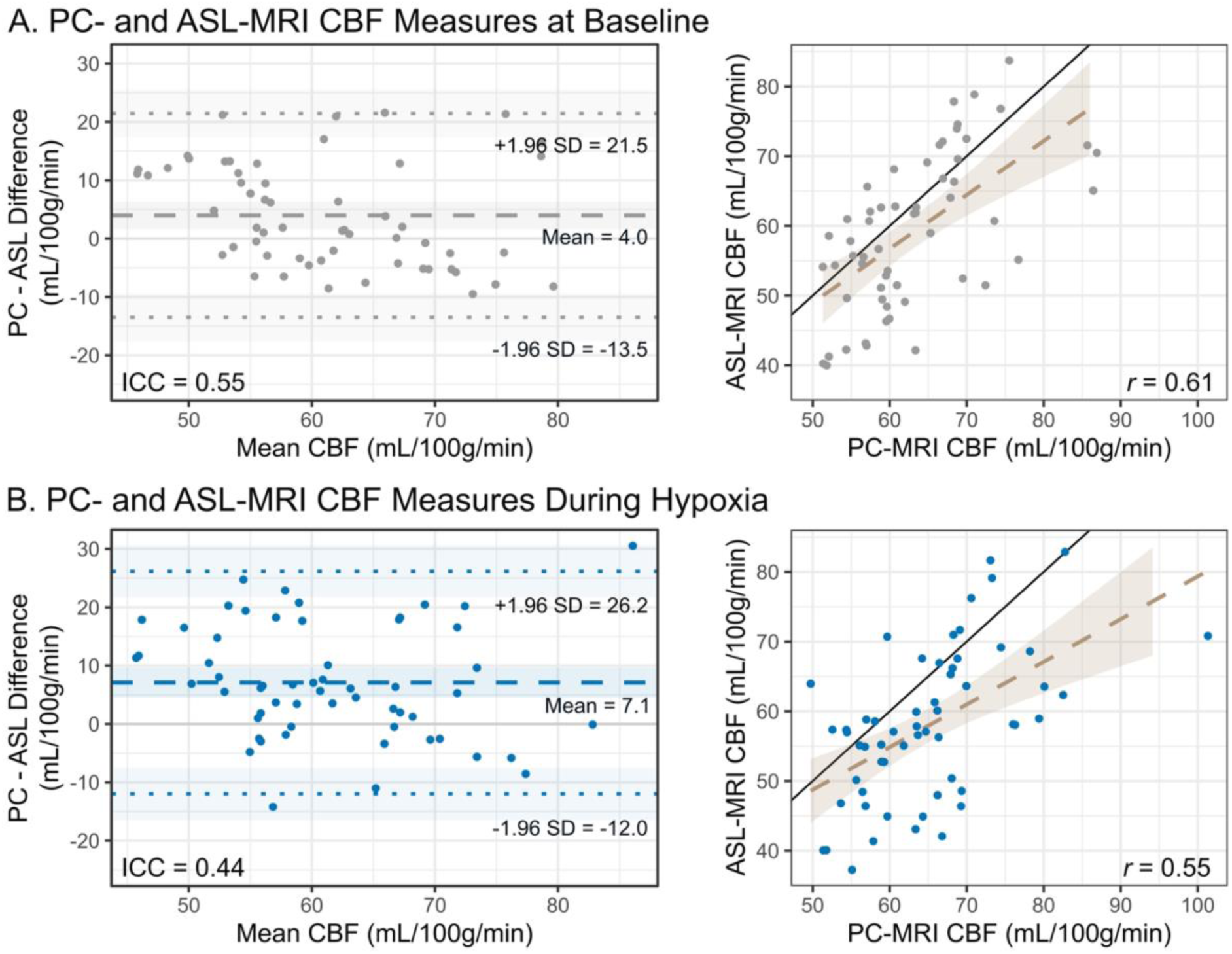
Phase Contrast and Arterial Spin Labeling Cerebral Blood Flow Measurement Agreement. Agreement between phase contrast (PC) and arterial spin labeling (ASL) measurements of whole-brain cerebral blood flow (CBF) and grey matter perfusion, respectively, are shown for measurements taken during baseline (A) and hypoxic (B) phases of scanning. Bland-Altman plots demonstrate the mean difference (dashed lines) and limits of agreement (dotted lines). Intraclass correlation coefficients (ICC, two-way mixed, absolute agreement) are also shown. Scatter plots demonstrate the divergence of the PC-ASL regression lines (dashed lines) from the identity line (solid black); Pearson’s correlation coefficients are included.

### 3.3 Blood Pressure

Acute intermittent hypoxia therapy did not significantly change blood pressure following a single session or over the three weeks of repeated sessions as compared to sham therapy (Table 1). While blood pressure values were significantly lower (p<0.001) at the end of intervention sessions compared to at the start of sessions with both sham and AIH treatments (Figure 4A), there was no significant difference in this effect between the two conditions (p=0.79 for MAP, p=0.42 for diastolic, p=0.53 for systolic). Thus, this reduction in blood pressure likely relates to the participant sitting quietly for half an hour in a laboratory environment rather than the gases being inhaled. Blood pressure was generally slightly reduced after 3 weeks of intervention exposure (Figure 4B), although this change was not significant for either sham or AIH treatment (p=0.10 for MAP, p=0.26 for diastolic, p=0.07 for systolic), and no significant difference between the two conditions for this effect was found (p=0.93 for MAP, p=0.45 for diastolic, p=0.24 for systolic). Individual changes in acute and longitudinal blood pressure measurements can be seen in Supplementary Figure 1.

**Figure 4.**
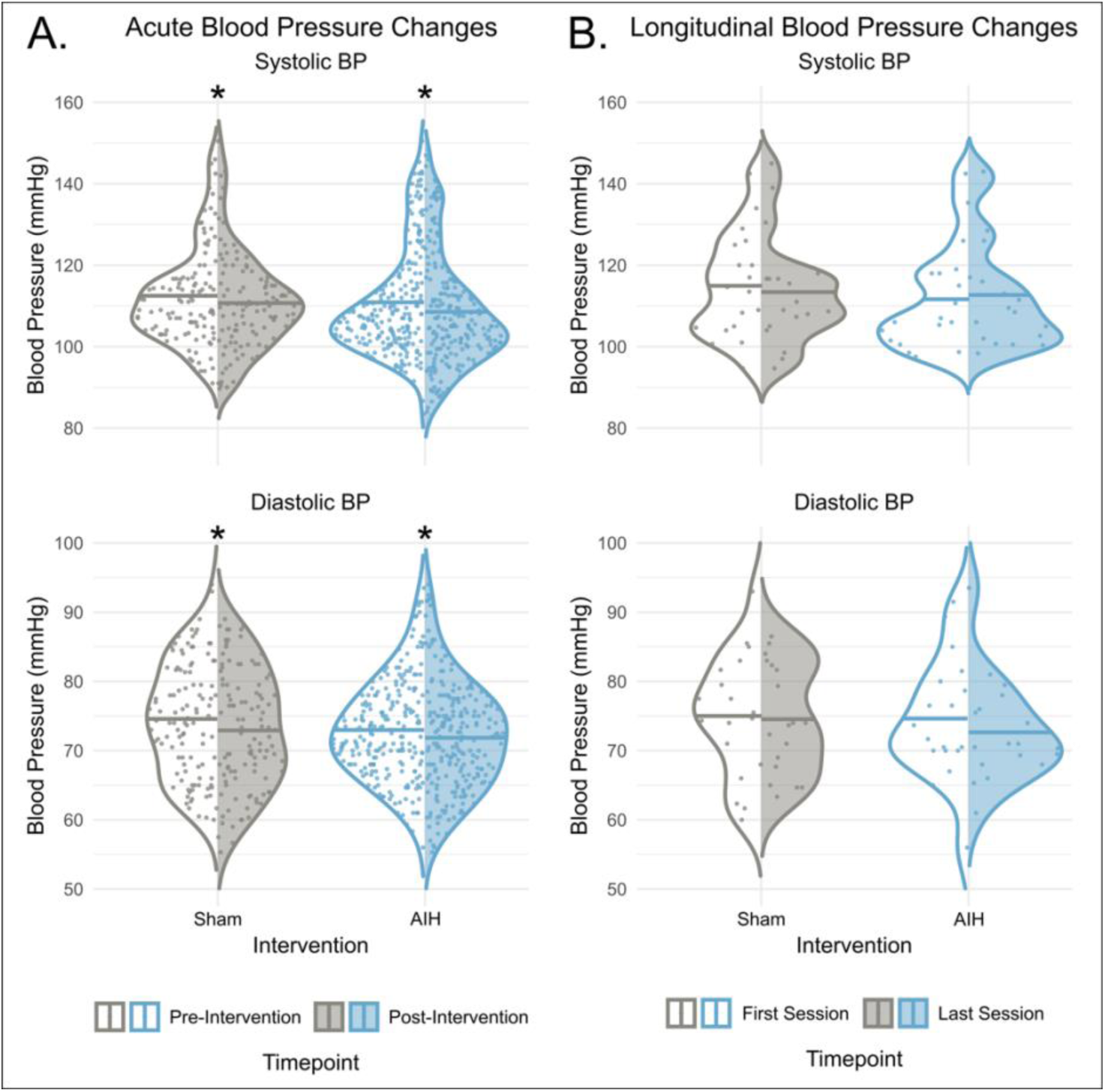
Changes in Blood Pressure with Intervention. A) Distributions of systolic and diastolic blood pressure measurements taken at the beginning and end of sham (grey) and acute intermittent hypoxia (AIH, blue) sessions. B) Distributions of systolic and diastolic blood pressure taken at the first and last intervention session for sham (grey) and AIH (blue) treatments. Significant differences, assessed with the Wilcoxon signed-rank test, are indicated by * (p<0.05, Bonferroni-corrected).

**Table 1.** Blood Pressure and Cerebrovascular Function Measures Before and After Intervention Arms for Sham and Acute Intermittent Hypoxia (AIH) Treatments. Values are listed as model-adjusted mean ± standard error. Effect magnitudes and p-values are reported for post-hoc pairwise comparisons, with * indicating p<0.05, Bonferroni-corrected. MAP: mean arterial pressure; CBF: cerebral blood flow; HX-CVR: cerebrovascular reactivity to hypoxia; PC: phase contrast MRI; ASL: arterial spin labeling MRI.

| Arm | Measure |  | Values |  | Effect Magnitude | p |
| --- | --- | --- | --- | --- | --- | --- |
|  |  |  | Before Intervention | After Intervention |  |  |
| Sham | MAP (mmHg) |  | 88.3 ± 2.13 | 87.5 ± 2.13 | 0.83 | 0.49 |
| | CBF ($\frac{mL}{100g}/min$) | PC | 64.1 ± 1.98 | 62.7 ± 1.98 | 1.47 | 0.41 |
|  |  | ASL | 62.3 ± 2.92 | 58.3 ± 2.92 | 4.00 | 0.10 |
| | HX-CVR<br>($-\frac{mL}{100g}/min$<br>% SpO<sub>2</sub>) | PC | 0.13 ± 0.04 | 0.04 ± 0.04 | 0.08 | 0.06 |
|  |  | ASL | -0.09 ± 0.05 | -0.01 ± 0.05 | 0.08 | 0.18 |
| AIH | MAP (mmHg) |  | 87.0 ± 2.13 | 86.0 ± 2.13 | 0.98 | 0.42 |
| | CBF ($\frac{mL}{100g}/min$) | PC | 61.9 ± 1.98 | 62.0 ± 1.98 | 0.08 | 0.97 |
|  |  | ASL | 58.0 ± 2.92 | 57.8 ± 2.92 | 0.43 | 0.90 |
| | HX-CVR<br>($-\frac{mL}{100g}/min$<br>% SpO<sub>2</sub>) | PC | 0.08 ± 0.04 | 0.06 ± 0.04 | 0.02 | 0.64 |
|  |  | ASL | 0.00 ± 0.05 | -0.03 ± 0.05 | 0.03 | 0.64 |

### 3.4 Cerebral Blood Flow

As shown in Table 1 and Figure 5, there was no significant change in whole-brain baseline CBF associated with AIH treatment compared to sham treatment as measured with PC MRI (p=0.60) or with pCASL MRI (p=0.42). Comparisons between the changes measured by the two modalities for each participant can be seen in Supplementary Figure 2. Furthermore, whole-brain CBF did not significantly change across the three-week intervention period with either treatment (p=0.64 for PC MRI, p=0.28 for pCASL MRI). The effect of the intervention did not change across vascular territories. Maps of group-averaged perfusion before and after each intervention are shown in Figure 6A, and group differences in perfusion are shown in Figure 6B with opacity modulated by difference t-statistics. Voxel-wise analyses revealed widespread decreases in perfusion in cortical gray matter with the sham intervention, whereas significant perfusion decreases were primarily localized to deep gray matter structures with the acute intermittent hypoxia intervention. When perfusion changes were compared between interventions (Figure 6C), significant increases associated with AIH treatment were found in the right frontal pole, right postcentral gyrus, and left precentral gyrus (interaction of time and experimental arm, p<0.05, uncorrected). Perfusion increases were also seen in the left postcentral gyrus and right precentral gyrus, suggesting a bilateral effect in the sensory-motor cortices, but these changes did not quite reach the significance threshold. Decreased perfusion was also seen in left frontomedial white matter, but this cluster is proximal to the left ventricle and likely an artifact of cerebrospinal fluid^60^. Differences in hypoxic burden had no significant effect on CBF (p=0.49 for phase contrast, p=0.31 for arterial spin labeling) or HX-CVR changes (p=0.63 for phase contrast, p=0.95 for arterial spin labeling).

**Figure 5.**
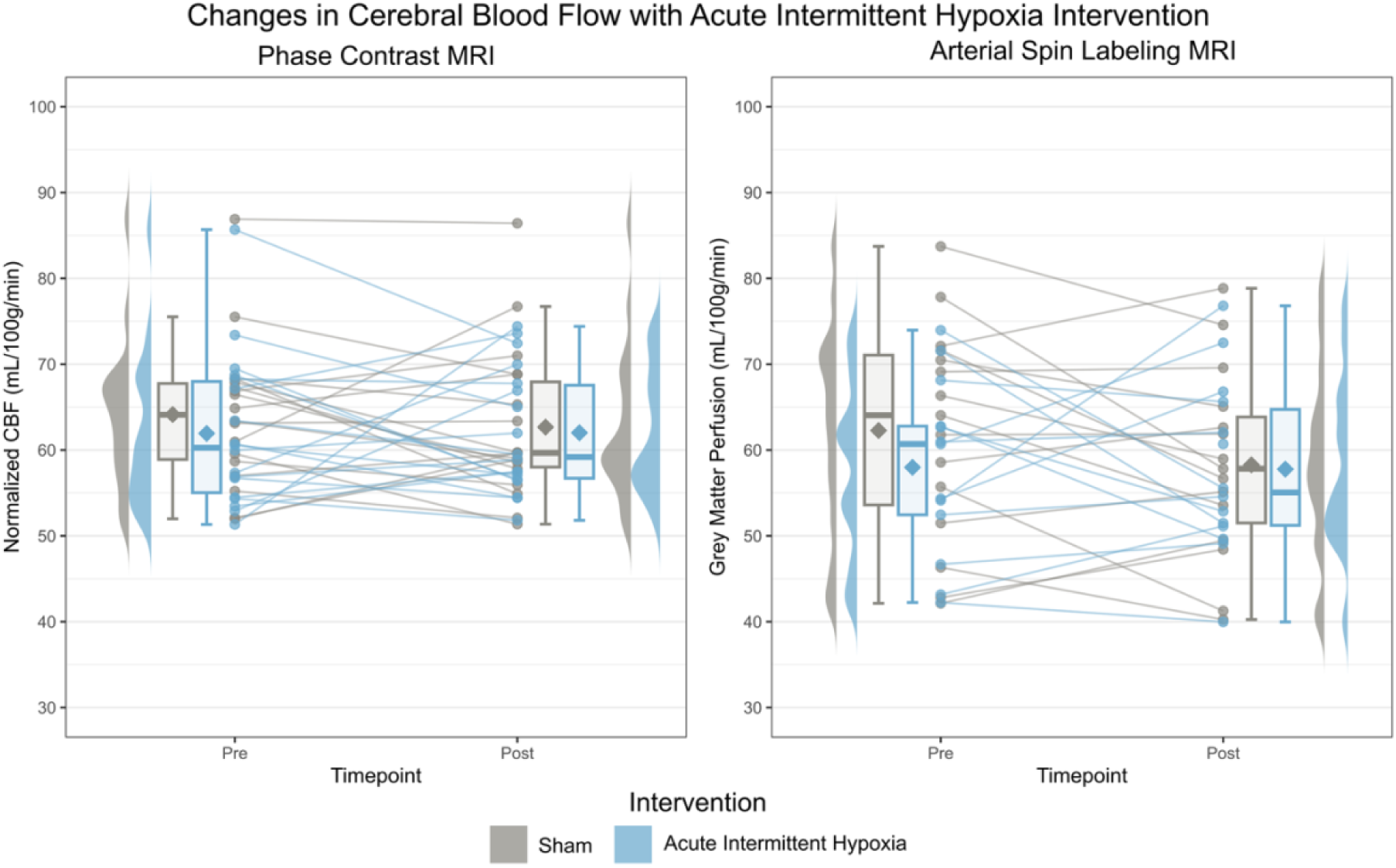
Changes in Whole-Brain Baseline Cerebral Blood Flow with Sham and Acute Intermittent Hypoxia Interventions. Changes in whole-brain resting cerebral blood flow (CBF), measured with phase contrast (left) and arterial spin labeling (right) MRI, are shown. Individual values measured before and after three weeks of sham (grey) or acute intermittent hypoxia (right) treatment are shown; some arterial spin labeling values were excluded due to data quality (15 out of 72). Ridge and box plots visualize data distributions, with diamonds indicating mean CBF for each timepoint and intervention.

**Figure 6.**
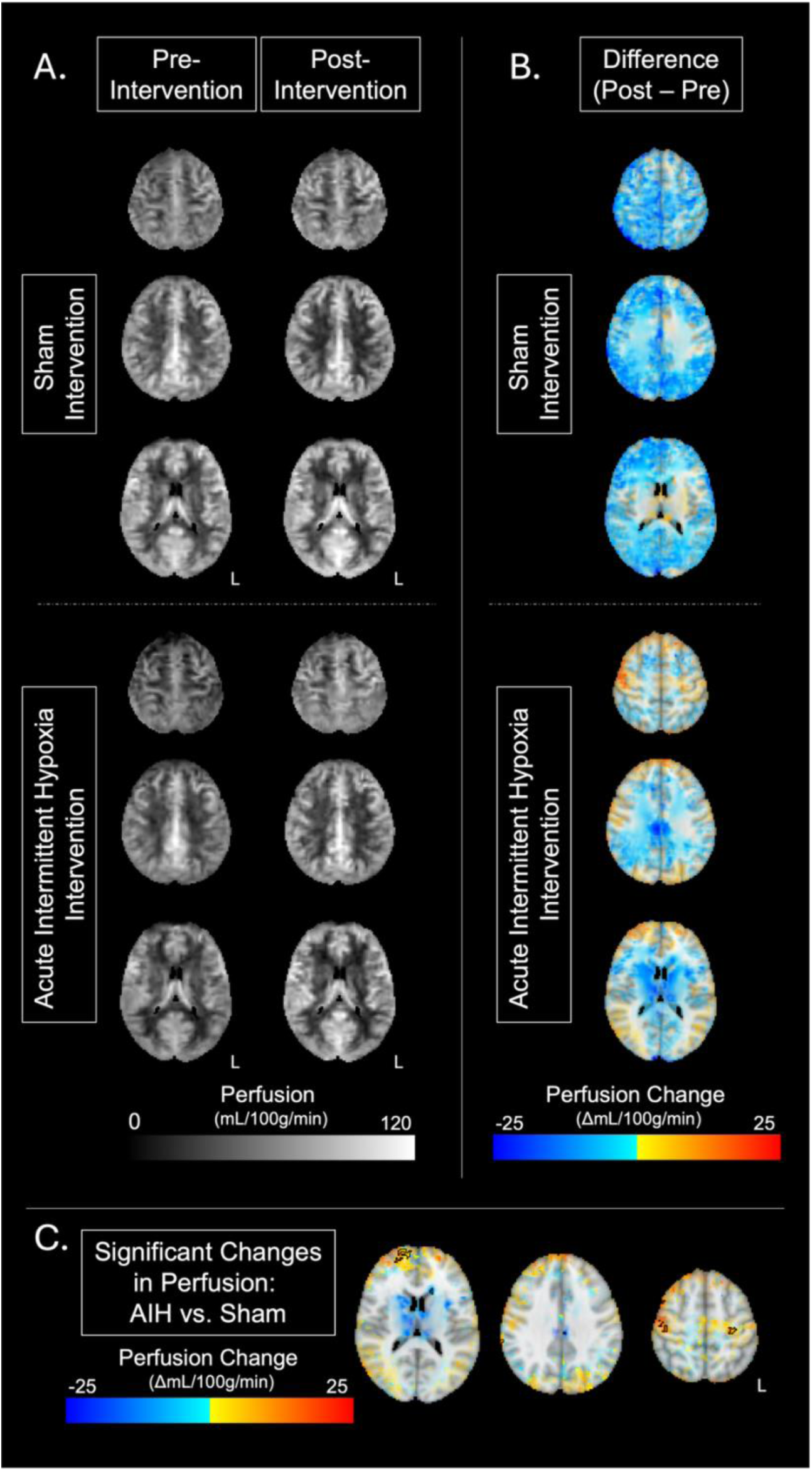
Group Maps of Perfusion and Perfusion Changes. Group-averaged perfusion maps are shown (A) before and after sham and acute intermittent hypoxia interventions. Overall group-averaged differences in perfusion (post - pre) are shown for each intervention (B), with the opacity of perfusion changes modulated by the difference t-statistics obtained from voxel-wise analyses of variance (p<0.05)^61^. Perfusion changes with the acute intermittent hypoxia (AIH) intervention as compared to the sham intervention are shown in panel (C), with significant clusters (n>40) outlined; significance for clusters and opacity modulation were determined using the interaction f-statistic (p<0.05, uncorrected). Increased perfusion was seen in the right frontal pole and bilateral sensory-motor cortices (pre- and postcentral gyri); identified clusters of decreased perfusion were proximal to ventricles.

### 3.5 Hypoxic Cerebrovascular Reactivity

The whole-brain cerebrovascular response to moderate sustained hypoxia, shown in Figure 7, did not change significantly with AIH or sham treatment, as measured by PC (p=0.31) or pCASL MRI (p=0.29). There was no significant change in HX-CVR between pre- and post-intervention MRI acquisitions (p=0.09 for PC, p=0.52 for pCASL). Of note, minimal and even negative blood flow responses to hypoxia were demonstrated in several sessions, at odds with the anticipated blood flow enhancement^62^. These paradoxical responses are seen in both PC and pCASL measures of HX-CVR and are largely consistent within individuals across time and measurement modality, although some instances of inconsistency between modalities were observed (shown in Supplementary Figure 2). While negative HX-CVR values were seen with both modalities, suggesting that this is an accurate measure of the response in some participants, the majority of pCASL measurements of HX-CVR were negative and may instead reflect methodological biases, such as reduced labeling efficiency in the pCASL data during enhanced arterial flow velocity^53^.

**Figure 7.**
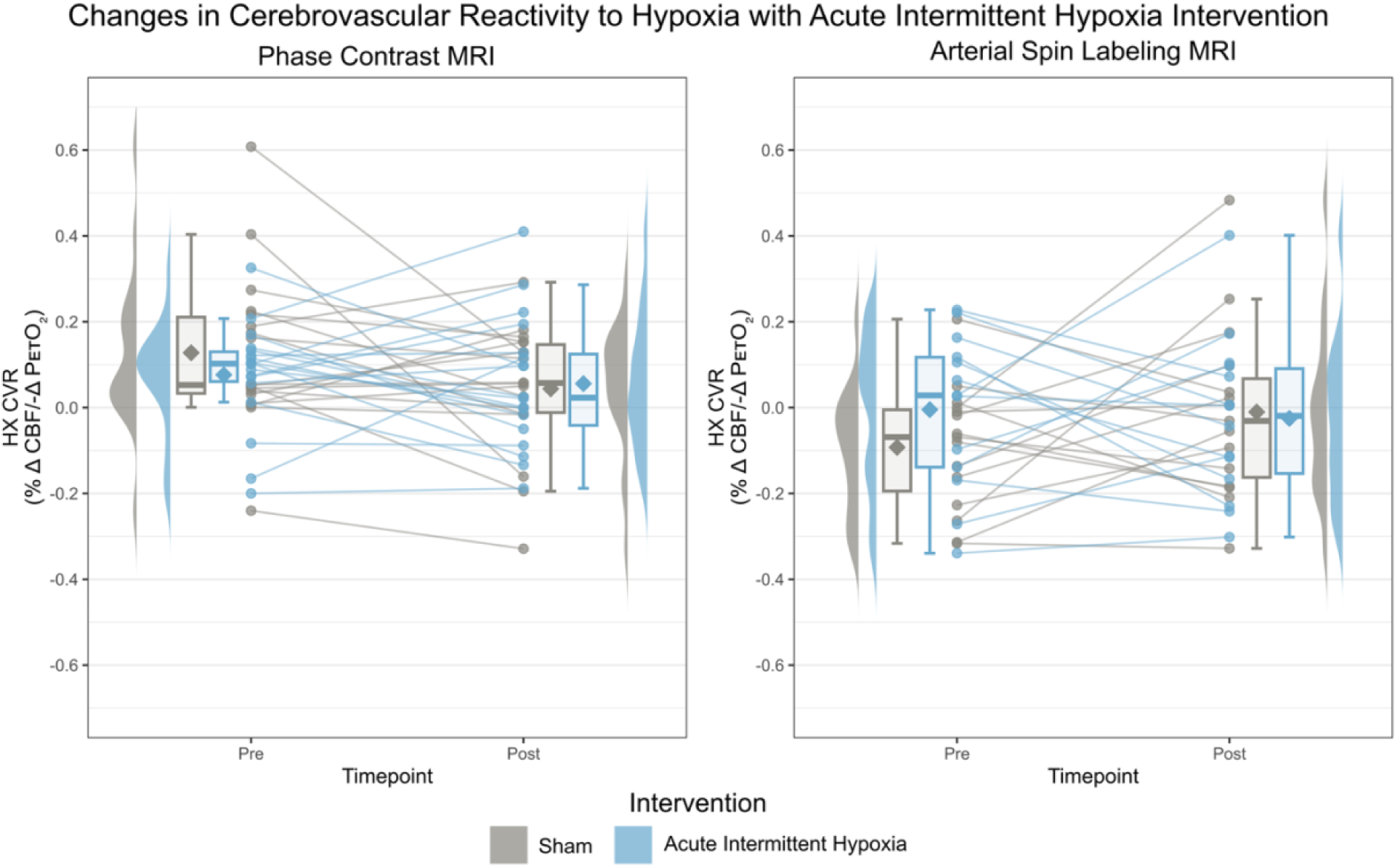
Changes in Hypoxic Cerebrovascular Reactivity with Sham and Acute Intermittent Hypoxia Interventions. Changes in hypoxic cerebrovascular reactivity (HX-CVR), measured with phase contrast (left) and arterial spin labeling (right) MRI, are shown. Individual values measured before and after three weeks of sham (grey) or acute intermittent hypoxia (blue) treatment are shown; some arterial spin labeling values were excluded due to data quality (15 out of 72). Ridge and box plots visualize data distributions, with diamonds indicating mean HX-CVR for each timepoint and intervention.

## 4 Discussion

This study sought to evaluate the impact of a three-week therapeutic AIH protocol on cerebrovascular function in young, healthy individuals. We found that this treatment protocol, delivered in 12 sessions over three weeks, did not significantly alter whole-brain CBF or HX-CVR as measured with phase contrast and arterial spin labeling MRI. Additionally, compared to sham therapy, this intervention did not significantly change blood pressure acutely or longitudinally. These results indicate that this therapeutic protocol did not induce sustained systemic changes in cerebrovascular function, supporting its potential safety in diverse clinical populations with varying cerebrovascular risk profiles. Focal increases in CBF were observed following AIH compared to sham, including in the pre- and post-central gyri (the primary motor and sensory cortices, respectively), and may reflect changes to the motor network, which has been implicated in previous studies of AIH.

### 4.1 Cerebrovascular Function Following Acute Intermittent Hypoxia Therapy

Whole-brain cerebrovascular function in healthy adults was not significantly impacted by this therapeutic AIH protocol. Cerebrovascular function following intermittent hypoxia has been previously studied in healthy adults with varying hypoxia protocols. These studies, assessing CBF and HX-CVR in healthy adults, have produced mixed results; the protocols and outcome measures of these studies are summarized in Table 2. In healthy adults, Zhang et al. found an 8% increase in middle cerebral artery velocity following AIH training, indicating enhanced CBF, whereas Foster et al. found that middle cerebral artery velocity did not change with AIH exposure^23,24^. Studies of HX-CVR following AIH training have similarly varied; Wang et al. and Zhang et al. found that HX-CVR generally increased after respective AIH protocols, contrasting with the reduced HX-CVR reported by Foster et al., as cerebrovascular resistance has a largely inverse relationship with CVR^23,26,27,29^. Animal studies of the effect of hypoxia on cerebrovascular function have also reported inconsistent results. A moderate AIH protocol (10 cycles of 5 minutes 13% oxygen alternated with 5 minutes normoxia daily for 14 days) elicited increased perfusion in mice whereas mixed effects have been reported with sustained hypoxia (7-12% oxygen sustained for 2-8 weeks), with some protocols inducing angiogenesis and others resulting in decreased CBF^63,64^. Notably, all previous studies of cerebrovascular function following AIH treatment in humans have used different AIH protocols (Table 2), and *all* produced a hypoxic dose more severe than the present study, typically utilizing hypoxic episodes of at least 5 minutes. The variability in results seen in previous studies of cerebrovascular function may be due to the variability between hypoxic protocols, which can substantially impact outcomes as discussed below. Variability within a protocol, however, does not appear to influence cerebrovascular outcomes, as differences in cumulative hypoxic burden did not significantly affect the measured outcomes in the present study. Variation in how the hypoxia dosage manifests in different individuals may not be a significant driver of cerebrovascular outcomes; rather, protocol timings and hypoxic targets may be more critical in determining the cerebrovascular effects of a protocol. It is therefore imperative to assess specific AIH protocols for their impacts on cerebrovascular function.

**Table 2.**
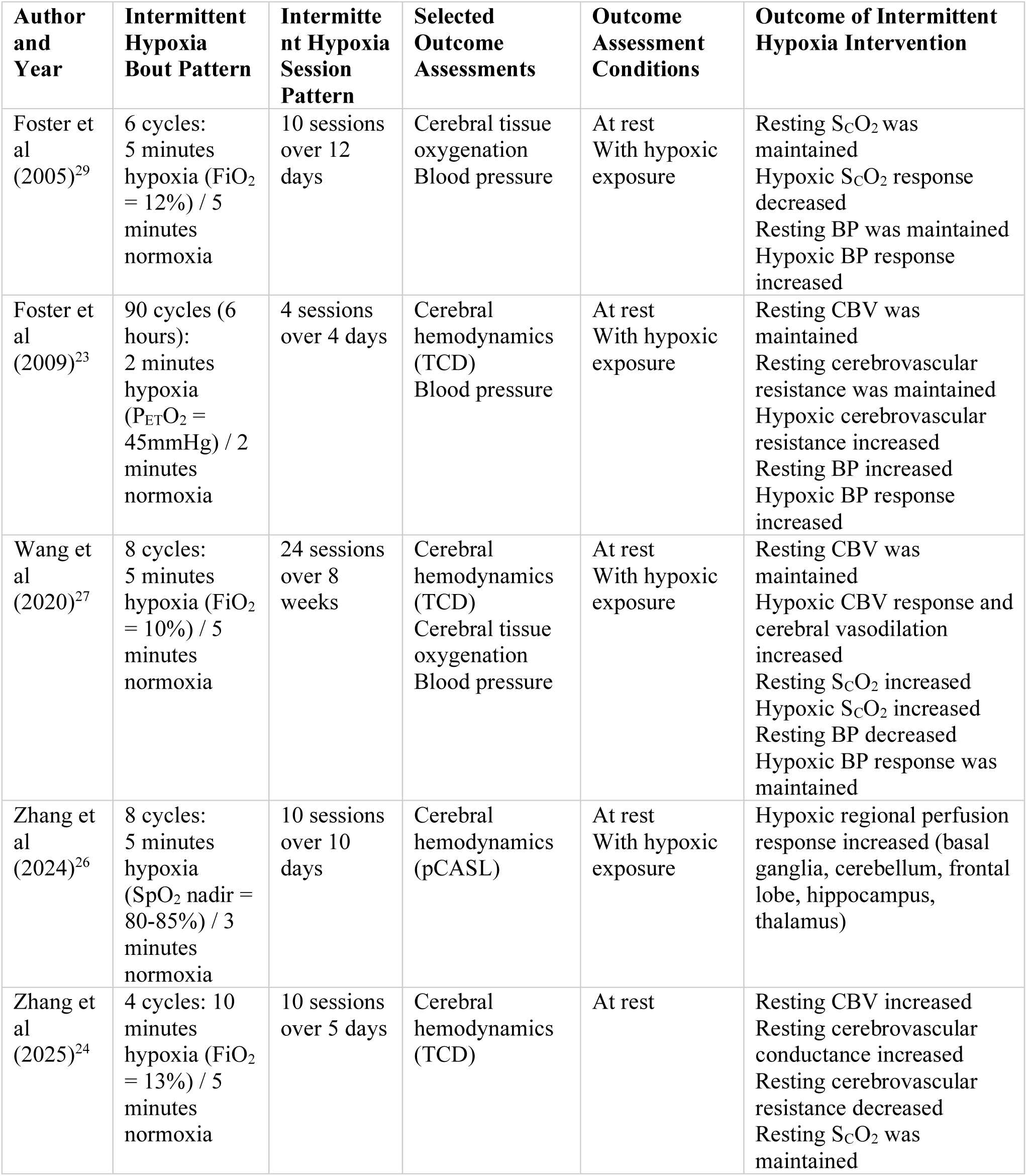
Summary of Intermittent Hypoxia Protocols and Outcomes in Studies of Cerebrovascular Function in Healthy Adults. CBV: cerebral blood velocity; FiO2: fraction of inspired oxygen; PETO2: partial pressure of end-tidal oxygen; SpO2: peripheral oxygen saturation; SCO2: cerebral tissue oxygenation; TCD: transcranial doppler ultrasonography.

| Author and Year | Intermittent Hypoxia Bout Pattern | Intermittent Hypoxia Session Pattern | Selected Outcome Assessments | Outcome Assessment Conditions | Outcome of Intermittent Hypoxia Intervention |
| --- | --- | --- | --- | --- | --- |
| Foster et al (2005)<sup>29</sup> | 6 cycles: 5 minutes hypoxia (FiO<sub>2</sub> = 12%) / 5 minutes normoxia | 10 sessions over 12 days | Cerebral tissue oxygenation<br>Blood pressure | At rest<br>With hypoxic exposure | Resting ScO<sub>2</sub> was maintained<br>Hypoxic ScO<sub>2</sub> response decreased<br>Resting BP was maintained<br>Hypoxic BP response increased |
| Foster et al (2009)<sup>23</sup> | 90 cycles (6 hours): 2 minutes hypoxia (P<sub>ET</sub>O<sub>2</sub> = 45mmHg) / 2 minutes normoxia | 4 sessions over 4 days | Cerebral hemodynamics (TCD)<br>Blood pressure | At rest<br>With hypoxic exposure | Resting CBV was maintained<br>Resting cerebrovascular resistance was maintained<br>Hypoxic cerebrovascular resistance increased<br>Resting BP increased<br>Hypoxic BP response increased |
| Wang et al (2020)<sup>27</sup> | 8 cycles: 5 minutes hypoxia (FiO<sub>2</sub> = 10%) / 5 minutes normoxia | 24 sessions over 8 weeks | Cerebral hemodynamics (TCD)<br>Cerebral tissue oxygenation<br>Blood pressure | At rest<br>With hypoxic exposure | Resting CBV was maintained<br>Hypoxic CBV response and cerebral vasodilation increased<br>Resting ScO<sub>2</sub> increased<br>Hypoxic ScO<sub>2</sub> increased<br>Resting BP decreased<br>Hypoxic BP response was maintained |
| Zhang et al (2024)<sup>26</sup> | 8 cycles: 5 minutes hypoxia (SpO<sub>2</sub> nadir = 80-85%) / 3 minutes normoxia | 10 sessions over 10 days | Cerebral hemodynamics (pCASL) | At rest<br>With hypoxic exposure | Hypoxic regional perfusion response increased (basal ganglia, cerebellum, frontal lobe, hippocampus, thalamus) |
| Zhang et al (2025)<sup>24</sup> | 4 cycles: 10 minutes hypoxia (FiO<sub>2</sub> = 13%) / 5 minutes normoxia | 10 sessions over 5 days | Cerebral hemodynamics (TCD) | At rest | Resting CBV increased<br>Resting cerebrovascular conductance increased<br>Resting cerebrovascular resistance decreased<br>Resting ScO<sub>2</sub> was maintained |

While overall measures of cerebrovascular function did not reveal changes with this AIH protocol, increases in regional perfusion in the frontal pole, postcentral gyri, and precentral gyri were seen with AIH in comparison to the sham intervention. Increased perfusion in these regions could indicate adaptations in cognition, sensory, and motor functions, respectively. In particular, cerebrovascular changes in the motor cortex may reflect alterations in motor network excitability, which has been extensively studied with similar therapeutic protocols in preclinical and human cohorts^31,65–67^. Previous research has found that repetitive AIH increased levels of vascular endothelial growth factor and brain-derived neurotrophic factor in the primary motor cortex of rats. The upregulation of these proteins promotes vascular remodeling and angiogenesis and is associated with neuroplasticity^65^. No association between cerebrovascular and motor changes was expected in the present study; further study of these effects in patient populations and their association with functional changes may establish mechanisms that link the cerebrovascular and motor changes that have been observed.

It is also important to acknowledge that AIH may have effects on cerebrovascular function beyond the changes in CBF and HX-CVR that were directly assessed in this study. This study did not incorporate measures of hematocrit, which typically displays an inverse relationship with CBF^68^. Decreases in CBF with prolonged hypoxia exposure, on the scale of days to years, are attributed to gradual increases in hematocrit, allowing for normal cerebral oxygen delivery despite low inhaled oxygen concentrations^69–72^. No intermediate measures of blood flow nor hematocrit were incorporated in the present study design across the three weeks, so it is possible that cerebrovascular function was initially enhanced by this AIH protocol but normalized over the three-week treatment period as hematocrit levels increased. However, previous studies of the oxygen-carrying capacity following AIH treatment suggest that this is not the case. Harding et al. found that four days of therapeutic AIH, delivered with a similar protocol to this study (4 sessions over 4 days, 15 bouts of 90 seconds per session), did not alter hemoglobin mass, whereas results from Tobin et al. indicate that 5 days of exposure to a more severe AIH protocol (5 sessions over 5 days, number and duration of bouts varied to sum to 70 or 72 minutes of hypoxic exposure per session) significantly increased hematocrit and red blood cell count^32,73^. Given the moderate level of AIH applied in the present study, significant hematological changes are unlikely.

Exposure to intermittent hypoxia forms reactive oxygen species (ROS) and may subsequently upregulate the expression of hypoxia-inducible factors (HIF) and vascular endothelial growth factors to stimulate angiogenesis and protection of the blood-brain barrier^74–76^. While ROS are required for beneficial neuroplasticity that is being targeted with AIH therapy and low, stable levels of ROS and HIF can benefit cerebrovascular and immune health, excess ROS and HIF produce inflammation and pathological cerebrovascular sequelae^77,78^. Direct measures of ROS and HIF serum values, and of angiogenesis and blood-brain barrier permeability, as have been incorporated in animal models and obstructive sleep apnea studies, could provide more conclusive insight into the effect on these cerebrovascular correlates^63,64,79^.

### 4.2 Influence of Acute Intermittent Hypoxia Protocol

The effects of therapeutic AIH are known to vary distinctly with the level of hypoxia, duration, and pattern of the exposure protocol^9^. The variations in protocol design and severity are likely responsible, in large part, for the disparity in findings in previous studies of the effect of AIH exposure on cerebrovascular function. The specific protocol implemented in this study was adapted from therapeutic practice in rehabilitation. Rehabilitative centers widely use a daily AIH stimulus consisting of 15 cycles alternating approximately 60 seconds of hypoxia (FiO2 9%) with room air^46,80–82^. This mild protocol has been found to elicit beneficial neuroplasticity without pathological effects^6,9^. Twelve AIH sessions were distributed across three weeks to reflect common rehabilitative therapy frequency, to maintain treatment frequency within the window established to produce neuroplasticity, and to reduce the practical burden on participants^83^. Protocols of similar severity, albeit abbreviated in their number of daily sessions, have demonstrated efficacy in enhancing motor function in chronic spinal cord injury, stroke, traumatic brain injury, and multiple sclerosis ^1,46,47,80–82,84,85^.

It is also understood that patient populations may be more susceptible to becoming hypoxic than healthy, able-bodied individuals when exposed to the same level of hypoxic air, as the SpO_2_ to FiO_2_ ratio is lower in many patient cohorts than in healthy individuals, likely due to enhanced chemosensitivity and blunted hypoxic ventilatory responses^86–89^. In this group of healthy adults, we saw great variability in the hypoxic burden, quantified as the integral of SpO_2_ desaturation below 90% over time (Figure 3), and low hypoxic burdens (<60%·s per bout) were frequent amongst this cohort. This protocol, while appropriate for patient populations as has been previously demonstrated, may have been insufficient to induce sustained physiological effects in healthy, able-bodied participants. Future work should build upon the foundation that this study lays to investigate cerebrovascular effects of this protocol in patient populations.

This study was limited in that our functional outcomes were only measured at one timepoint following treatment arms. Imaging was performed one or three days after the final intervention session, thus reflecting both cumulative and acute effects; acute cerebrovascular effects resulting from AIH treatment may have been underestimated due to this delay, as such effects may have been more apparent in immediately after the intervention. However, studies of the motor effects of therapeutic AIH have indicated that, while limb function changes peak 3 hours after the intervention, they were still present more than 24 hours later. Furthermore, preclinical research has shown that cerebrovascular changes persisted for more than 6 weeks following prolonged chronic exposure to moderate hypoxia^64^. If significant changes in systemic cerebrovascular function are in fact present with this protocol, they are not sustained.

### 4.3 Magnetic Resonance Imaging Measures of Cerebrovascular Function

Phase contrast and ASL are both commonly used to measure CBF, but as evidenced by this study, these quantifications are not interchangeable, and we observed substantial fixed and proportional biases in both resting and hypoxic CBF measurements. CBF was consistently underestimated by ASL as compared to PC measures, whereas grey matter CBF values are expected to be greater than whole-brain blood flow^90^. ASL measurements, in particular, are limited in their reliability as the perfusion calculations are reliant on a number of assumptions, including consistent labeling efficiency and uniform oxygenation and hematocrit (and therefore blood T1)^49^. The observed fixed offsets suggest that the labeling efficiency value used (α=0.8) may overestimate the true labeling efficiency achieved; this is supported by the higher fixed bias seen under hypoxia, when blood velocity generally increases and labeling efficiency should therefore decrease^53,91^. This higher fixed bias under hypoxia could additionally result from changes in the T1 of blood, as the T1 shortens with blood deoxygenation^92^. The proportional bias indicates PC overestimation or ASL underestimation at low flow, which may be due to inappropriate velocity encoding or arterial transit time elongations, respectively, among other contributors.

The variability in CBF and CVR measurements has substantial implications for future studies assessing the effects of a hypoxia-based intervention or stimulus on cerebrovascular function. Overall variability in baseline and moderate hypoxic measures of CBF was in line with previous literature, which reported total coefficients of variation of up to 25% for PC CBF measurements across a similar time frame^93^, but we saw within-subject variance that exceeded between-subject variance, suggesting poor-to-moderate longitudinal reliability of this measure. Based on these estimates of variability in PC data, a greater sample size (n=34) would be needed to detect a moderate increase (Cohen’s d = 0.5) in whole-brain baseline CBF with the same experimental design.

HX-CVR, in particular, showed poor repeatability, as explored in Johnson et al^62^. We also observed negative HX-CVR in a small number of PC datasets and a larger number of ASL datasets. Although negative HX-CVR measured with ASL may reflect underestimation of blood flow estimates due to reduced labeling efficiency during hypoxia, PC estimates of CBF theoretically do not share this confound. It remains an open question as to why certain individuals demonstrate absent or paradoxical *decreases* in CBF during a sustained hypoxia stimulus. A larger sample size would allow stratification by participant characteristics to examine which baseline physiological or demographic factors may explain this inter-individual variability in response to hypoxia. However, avoiding recruitment bias is challenging given the need to prioritize safety in line with research study inclusion and exclusion criteria. There is also a potential bias arising from differences in participants’ ability to complete all experimental sessions to sufficient data quality standards and while maintaining safety.

## 5 Conclusion

In this study, we have shown that this clinically relevant AIH protocol imposes no sustained systemic cerebrovascular changes in a cohort of healthy younger (∼30yo) adults. This work paves the way for expanding access to a promising therapeutic intervention and lays the foundation for assessment of the impact of AIH therapy on cerebrovascular function in clinical populations. Furthermore, we observed that an AIH protocol relevant to therapeutic motor rehabilitation significantly increased focal perfusion in select cortical regions, including primary motor cortices. The motor cortex has been implicated in previous studies examining the effects of this therapeutic intervention on respiratory and non-respiratory motor function, and the perfusion changes observed here may reflect these motor function changes. Finally, the results of this work suggest that improved reliability of imaging metrics and controlling for variability in AIH protocols delivered in clinical studies would allow us to better characterize true variability in the physiological response to AIH therapy, expanding our understanding of patient populations in which this rehabilitative therapy could be beneficial.

## 7 Author Contributions

HJ: Investigation, Formal analysis, Project administration, Visualization, Writing – original draft, Writing – review & editing. MW: Investigation, Writing – review & editing. RS: Conceptualization, Methodology, Investigation, Writing – review & editing. YC: Conceptualization, Methodology, Writing – review & editing, Funding Acquisition. TP: Conceptualization, Methodology, Writing – review & editing, Funding Acquisition. FS: Conceptualization, Methodology, Writing – review & editing, Funding Acquisition. MB: Conceptualization, Methodology, Project Administration, Writing – review & editing, Supervision, Funding Acquisition.

## 9 Acknowledgments

The authors would like to thank Rachael Young and the Center for Translational Imaging at Northwestern University for supporting data collection efforts, as well as Milap Sandhu and colleagues at the Shirley Ryan AbilityLab for ongoing exchange of best practices in AIH delivery. s

