## Supplementary for "Therapeutic acute intermittent hypoxia does not alter whole-brain cerebrovascular function in healthy individuals"

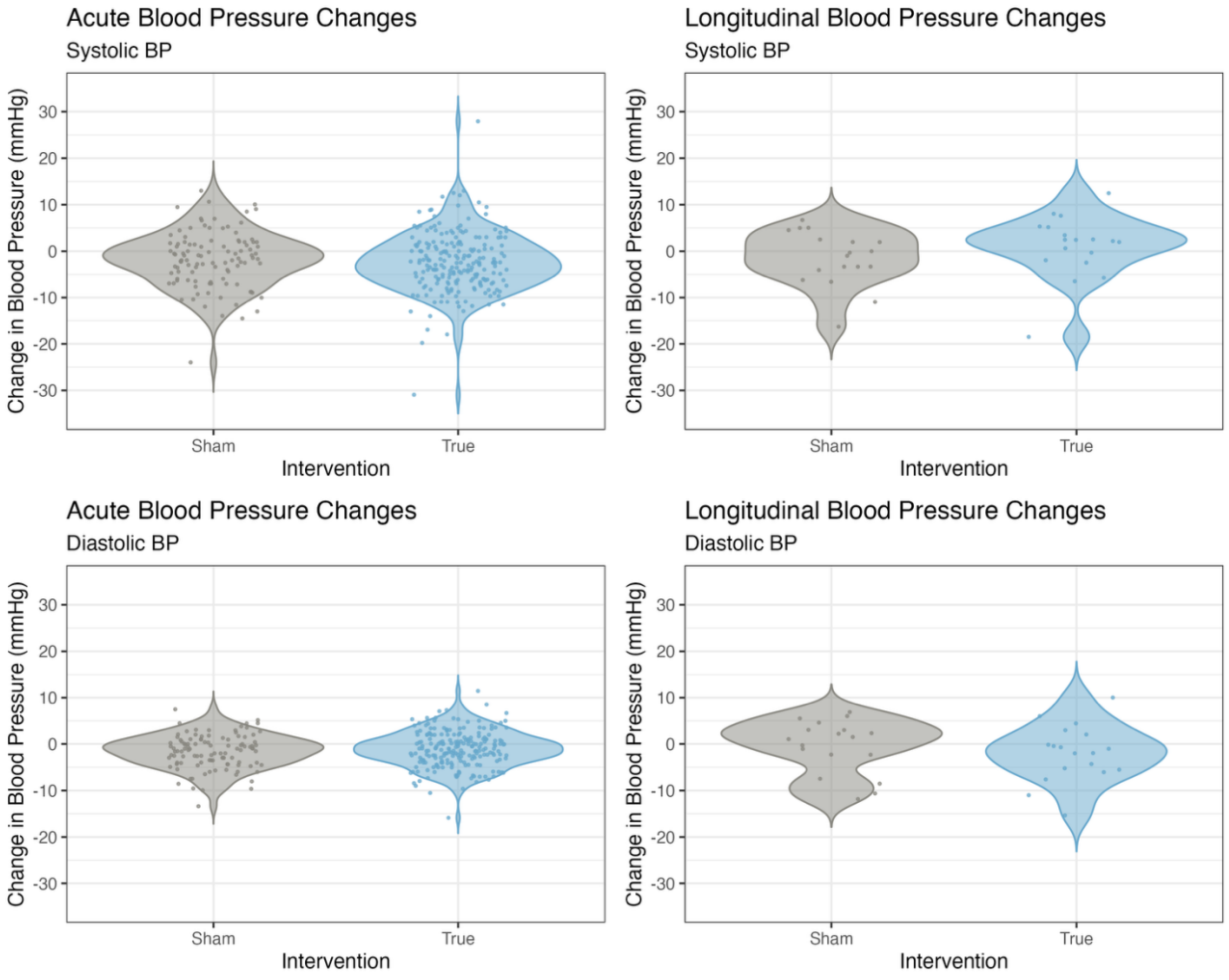


*Supplementary Figure 1. Blood Pressure Differences with Intervention.* A) Distributions of differences in systolic and diastolic blood pressure measurements across sham (grey) and acute intermittent hypoxia (AIH, blue) sessions, with the pre-session average value subtracted from the post-session average value. B) Distributions of differences in systolic and diastolic blood pressure across the intervention period for sham (grey) and AIH (blue) treatments, with the first-session average value subtracted from the last-session average value.

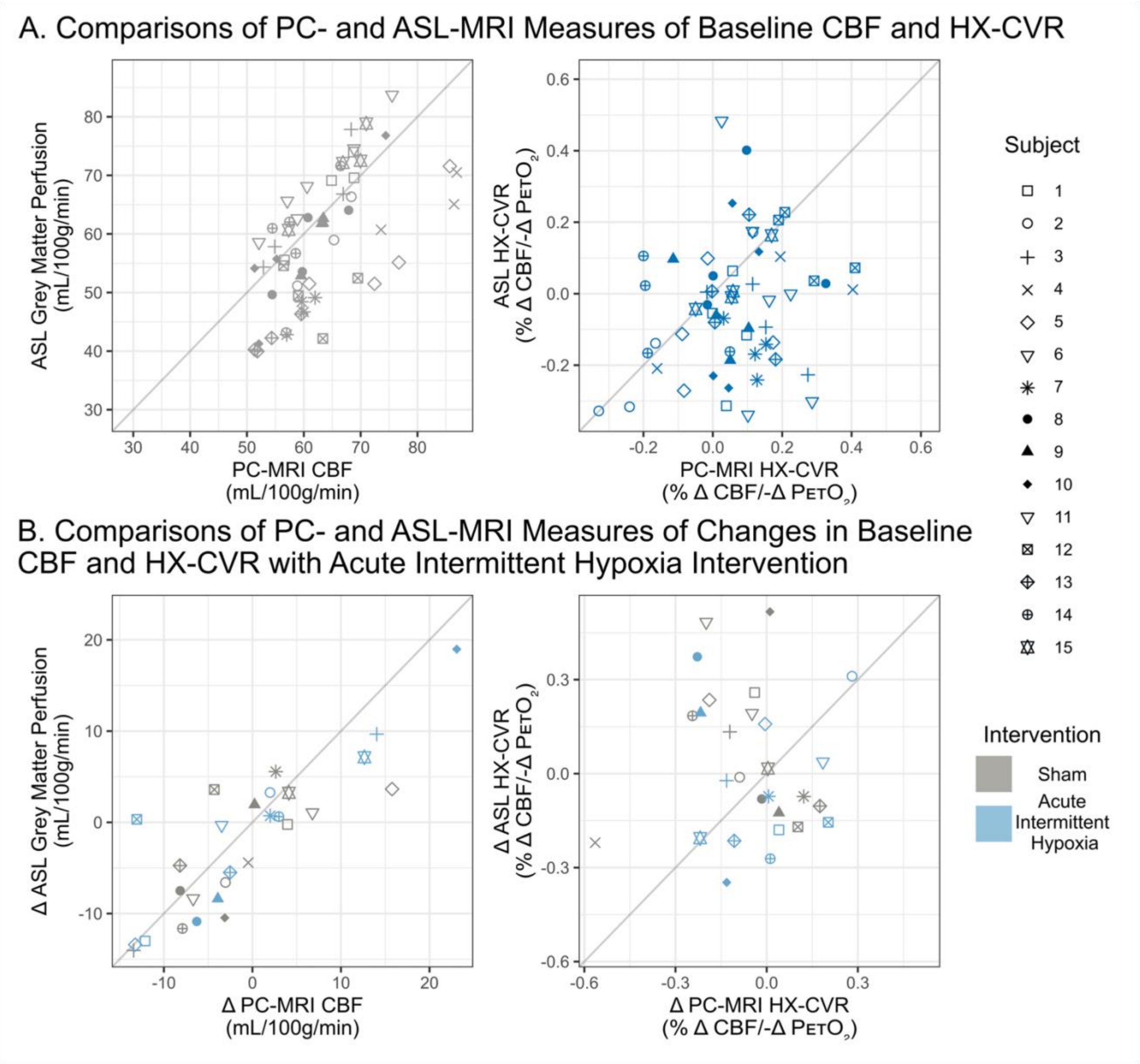


*Supplementary Figure 2. Subject-coded Baseline CBF and HX-CVR and Changes with Sham and AIH Interventions compared between PC- and ASL-MRI.* Comparisons of measures of cerebral blood flow (CBF) during baseline respiratory conditions (left) and measures of hypoxic cerebrovascular reactivity (HX-CVR; right) as obtained with phase contrast (PC) and arterial spin labeling (ASL) magnetic resonance imaging (MRI) are shown in Panel A. Individual subjects are represented by different shapes. Comparisons of the change across the intervention period (post - pre) in baseline CBF (left) and in HX-CVR (right) as measured by PC- and ASL-MRI are shown in Panel B; the sham intervention is shown in grey, while the acute intermittent hypoxia intervention is shown in blue. Note that some data points are missing as measures from ASL-MRI were excluded due to data quality.
